# Computational Assessment of the Regulation-Modulating Potential for Noncoding Variants

**DOI:** 10.1101/819409

**Authors:** Fang-Yuan Shi, Yu Wang, Dong Huang, Yu Liang, Nan Liang, Xiao-Wei Chen, Ge Gao

**Author notes:** Email: FYS, YW, DH, YL, NL, XWC, GG.

## Abstract

Large-scale genome-wide association and expression quantitative trait loci studies have identified multiple noncoding variants associated with genetic diseases via affecting gene expression. However, effectively and efficiently pinpointing causal variants remains a serious challenge. Here, we developed CARMEN, a novel algorithm to identify functional noncoding expression-modulating variants. Multiple evaluations demonstrated CARMEN’s superior performance over state-of-the-art tools. Its higher sensitivity and low false discovery rate enable CARMEN to identify multiple causal expression-modulating variants that other tools simply missed. Meanwhile, benefitting from extensive annotations generated, CARMEN provides mechanism hints on predicted expression-modulating variants, enabling effectively characterizing functional variants involved in gene expression and disease-related phenotypes. CARMEN scales well with the massive datasets and is available online as a Web server at http://carmen.gao-lab.org.

---

Approximately 98% of the human genome is noncoding^1^. While genome-wide association studies (GWAS) have identified thousands of disease-associated or complex trait-associated variants^2, 3^ and more than 90% of disease-associated variants are located in noncoding regions^4^, their biological functions and mechanisms remain elusive^5, 6^.

Several tools have been proposed to prioritize functional noncoding variants based on existing annotations at corresponding loci^7–10^. Recently, convolutional neural networks (CNNs)^11^ have been widely used to characterize the regulatory activity of sequences, enabling the calling of regulatory variants that result in significant changes in binding affinity^12, 13^ based on sequence only.

The massively parallel reporter assay (MPRA) enables the systematic screening of potential regulatory variants and provides valuable candidate training datasets to generate *in silico* prediction tools that predict the variants’ effects on gene expression. EnsembleExpr^14^ uses MPRA data to predict the causal variants in eQTLs, and ExPecto^15^ *ab initio* predicts the variants’ effects on gene expression from 40-kb promoter-proximal sequences based on reference data but not mutagenesis data.

Here, we developed CARMEN, an algorithm framework for predicting the effects of noncoding variants on both gene expression and disease risk (Fig. 1a). Compared with state-of-the-art tools, CARMEN shows superior performance on both high-throughput datasets and low-throughput case studies and has higher sensitivity with low false discovery rate, enabling effective identification of multiple causal expression-modulating variants that other tools missed. We also illustrated that CARMEN can pinpoint causal variants other than the reported lead SNPs in GWAS and eQTLs. CARMEN is available as a Web server with free access for academic usage at http://carmen.gao-lab.org.

**Figure 1:**
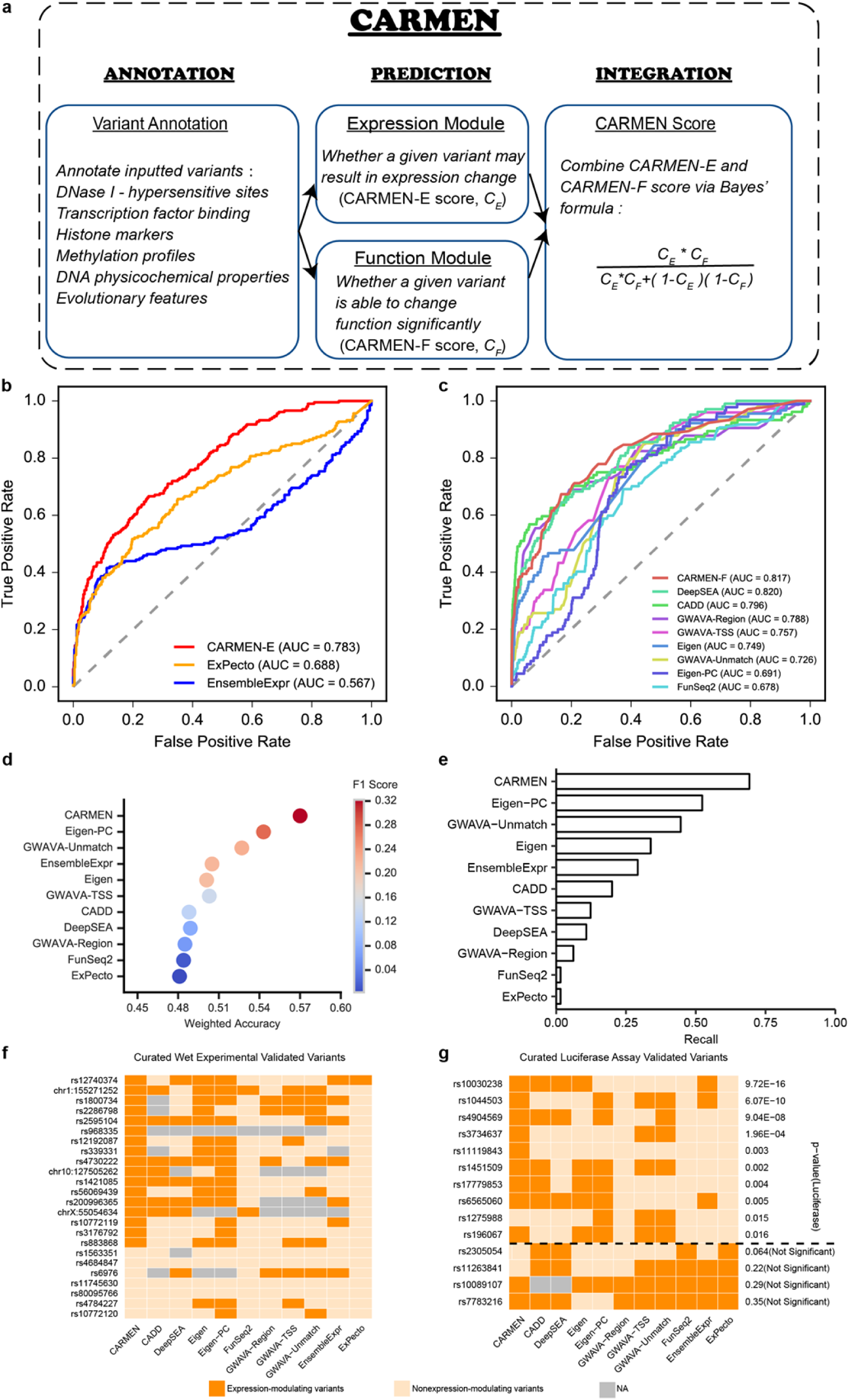
**a**, Overview of the CARMEN workflow. **b**, The ROC curve for the CARMEN-E module. The performance was evaluated on the testing dataset of the curated MPRA dataset with 738 variants (207 positive variants, 531 control variants). The ExPecto predicted scores were calculated as the average absolute score of 208 tissues. **c**, The ROC curve for the CARMEN-F module. The performance was evaluated on a pathogenic dataset with 2,135 variants (Supplementary Table 7). **d**, The performance comparison of CARMEN to other tools^26^ bubbles colored by F1 score. The thresholds of the different tools were obtained from respective official websites or reference papers (Supplementary Table 10). **e**, The recall of different tools on the known expression-modulating variants^26^. The X axis indicates the proportion of variants from the significant expression-modulating variants that were predicted as positive variants by the different tools. **f**, Performance comparison on a curated list of experimentally characterized variants validated by various low-throughput technologies from the literature (Supplementary Table 3). **g**, Performance comparison on an independent luciferase validated dataset^60^ (Supplementary Table 3). The right y-axis represents the reported *p* values in the luciferase assays; the four below the dotted line are the negative variants. Of note, the birth-weight-associated variant rs11119843^61^ is missed by all other tools.

## Results

Multiple studies demonstrate that noncoding variants can modulate gene expression via several distinct mechanisms^16–18^. Both TF binding and epigenetic modifications contribute to gene regulation^19, 20^. Inspired by DeepSEA^12^, we trained CNN models with large-scale chromatin-profiling data generated by Encyclopedia of DNA Elements (ENCODE) and Roadmap Epigenomic Project directly, including 1,249 TF binding profiles for 596 distinct TFs, 280 DHS profiles, 766 histone-mark profiles and 108 DNA methylation profiles assessed in 36 cell lines and tissues (see Methods for more details as well as Supplementary Fig. 1 and Supplementary Table 1). Networks were trained and tuned for each profile separately, resulting in 2,403 highly accurate models with an average AUROC (Area Under the Receiver Operating Characteristic Curve) of 0.908 and an average AUPRC (Area Under the Precision Recall Curve) of 0.904 (Supplementary Fig. 2b and Supplementary Table 2). Compared to DeepSEA, CARMEN not only covers more than twice as many features (Supplementary Fig. 2a) but also shows superior accuracy (Supplementary Fig. 3, median AUROC 0.973 vs. 0.935, single-tailed Wilcoxon-test *p* = 1.747 × 10^−25^; median AUPRC 0.975 vs. 0.357, single-tailed Wilcoxon-test *p* = 5.256 × 10^−154^).

DNA physicochemical properties also affect gene expression through influencing the shape properties of flanking sequences near transcription factor binding sites^21^. We incorporated 13 conformational and physicochemical DNA properties^22^ to evaluate the noncoding variants’ effects on DNA shape change.

Finally, along with the eight PhyloP^23^ and PhastCons^24^ conservation scores derived from multiple genome alignments in the primate/mammal/vertebrate clades, CARMEN generated 2,424 candidate features for each inputted variant.

To reduce the risk of overfitting and high time costs of model training, we adopted a data-driven feature selection approach before model training. Briefly, we pretrained a multilayer neural network and estimated features contributions by comparing the difference in the activation value of each neuron to its ‘reference activation’^25^. Using the distribution of contribution scores from each dataset, we selected the features with absolute contribution scores greater than the threshold (Supplementary Fig. 6) for follow-up training (**Methods**).

To evaluate the gene expression-modulating potential of a given variant, we trained a dedicated module, CARMEN-E, based on a manually curated MPRA dataset (**Methods**). Trained with 689 features choice from the data-driven feature selection approach, the CARMEN-E module had significantly superior performance than both EnsembleExpr and ExPecto (Fig. 1b). Additionally, a separate module, CARMEN-F, was trained on the HGMD Disease-causing Mutation (DM) in a regulatory-variant dataset to identify disease-causing variants with high accuracy (cross-validation AUROC = 0.921, also see Fig. 1c for CARMEN-F evaluation on an independent pathogenic dataset).

To characterize the disease-causing variants that function through modulating gene expression, the outputs of CARMEN-E and CARMEN-F were further integrated as the CARMEN score:

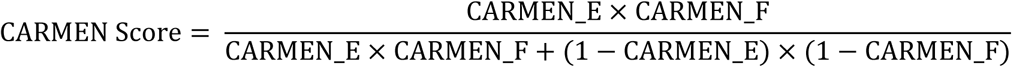

which will be used in the following evaluations.

### CARMEN shows superior performance on both large-scale independent datasets and experimentally characterized loci than state-of-the-art tools

To validate the robustness of CARMEN, we first tested it on two independent datasets generated by STARR-seq^26^.

The cancer-risk dataset^26^ consists of 1,164 curated regulatory positive and 5,375 control variants (see Methods for more details on data curation as well as Supplementary Fig. 7a). Given its significantly unbalanced nature, we employed the F1 score and weighted accuracy for performance comparison. CARMEN shows significantly superior performance than the state-of-the-art tools^7, 10, 12, 14, 15, 27, 28^. Of note, we found that DeepSEA and ExPecto performed worse, partly due to their surprisingly low sensitivity (Fig. 1d). Further inspection of allele-specific expression (ASE) variants showed that CARMEN successfully called 69.2% (45 out of 65) of significant allele-specific expression-modulating variants, while DeepSEA and ExPecto found only 7 and 1, respectively (Fig. 1e, also see Supplementary Fig. 7b, c, d for evaluation on the additional dataset).

We further compared the performance on a curated list of experimentally characterized variants validated by various low-throughput technologies from the literature^29, 30, 31–38, 39–41^. Out of 24 experimentally validated variants, 17 were correctly reported as positive by CARMEN, showing the highest sensitivity among others (Fig. 1f and Supplementary Table 3).

Notably, benefitting from the extensive annotation generated, CARMEN is able to provide hints on possible mechanisms for how the variant changes the gene expression. For example, the variant rs883868 has been shown to modulate the expression of UBASH3A by disrupted binding of transcription factor YY1^41^, which is consistent with the output of the CARMEN annotations (Fig. 2).

**Figure 2:**
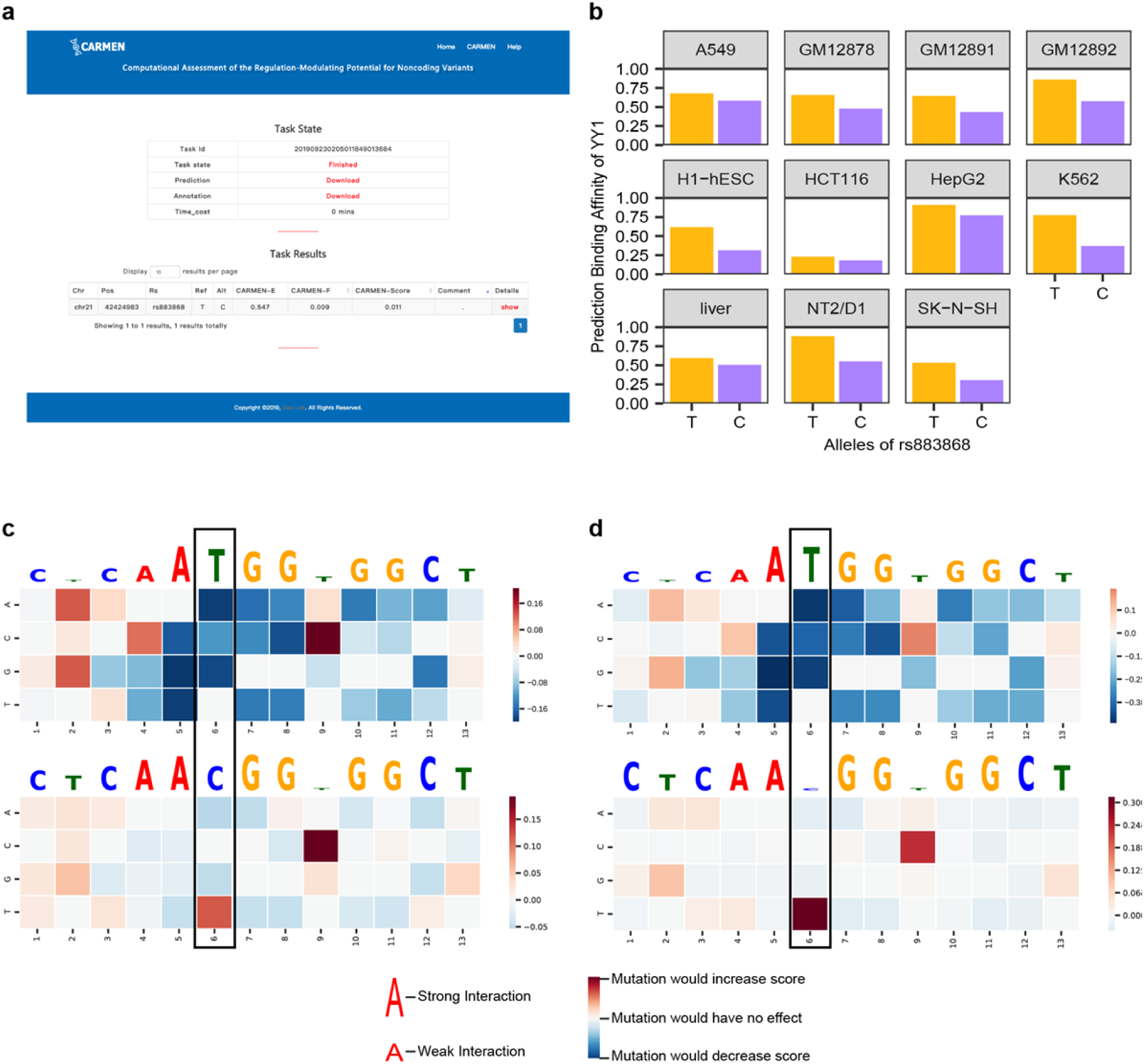
CARMEN helps pinpoint the functional mechanism of noncoding expression-modulating variant. **a**, CARMEN predicted rs883868 as expression-modulating variant which was further validated by independent CRISPR/Cas experiment^41^. **b**, The CARMEN annotation indicated that the alternative allele C decreases the binding affinity of YY1, which was also validated via 3C-qPCR^41^. **c and d**, Mutation maps for the variant effects on YY1 transcription factor binding in GM12878 (**c**) and K562 (**d**).

### CARMEN can pinpoint causal variants other than the GWAS-reported lead SNPs

Large-scale association studies, such as GWAS and eQTLs, have identified a number of genetic variants associated with complex human diseases and traits. However, a gap between the association of a locus and the causal variant still exists because many inherited variants and sentinel variants are in strong linkage disequilibrium regions^3, 5, 42^. We applied CARMEN on 51,878 reported lead SNPs extracted from the GWAS Catalog (https://www.ebi.ac.uk/gwas/home) and found that variants reported by multiple studies obtained significantly higher CARMEN scores than those that were not (single-tailed Wilcoxon-test *p* = 4.233 × 10^−39^), confirming previous observations^27^.

To pinpoint potential causal variants other than lead SNPs, we ran CARMEN on both the reported lead SNPs and variants with strong LD (r^2^ >=0.75) and found that 45.33% of the reported lead SNPs show significantly weaker regulatory potential than nearby variants within the same LD block (r^2^ > 0.75). While most of the differences were modest, 6.65% of variants showed differences larger than thirty-fold. (Supplementary Fig. 8)

For example, several GWA studies report that variant rs1701704 is associated with the susceptibility of type 1 diabetes (*p* < 5 × 10^−8^)^43–45^, but CARMEN found that this variant shows only very weak regulatory potential (CARMEN score = 0.0024). Meanwhile, CARMEN pinpointed a nearby variant rs705698 with rather high potential (CARMEN score = 0.4070, Fig. 3a). While rs705698 has few annotations in the UCSC Genome Browser^46^ (Supplementary Fig. 10a,b), the variant falls into the first intron of gene RAB5B, which is a candidate gene of type 1 diabetes^44^ with strong linkage (r^2^=0.90746), and the annotation module of CARMEN suggests that it is a conserved locus with YY1 changed in the K562, HepG2, GM12892 cell lines, YY2 changed HEK293 cell line, CBX5 and CBX1 changed in the K562 cell line (Supplementary Fig. 11). Independent luciferase assays further confirmed the significant change in report expression for rs705698 but not for rs1701704 (Fig. 3b and Supplementary Table 11). Notably, while CARMEN presents crystal clear contrast on the lead SNP and causal SNP, many state-of-the-art tools missed them (Fig. 3e and Supplementary Table 9). Likewise, variant rs1727313 was reported as an SNP associated with type 2 diabetes (*p* = 1 × 10^−8^)^47^. CARMEN found no regulatory potential for this variant (CARMEN score = 0), but the linkage variant rs146239222 showed high regulatory potential (CARMEN score = 0.1999, Fig. 3c), which was consistent with our luciferase reporter assay (Fig. 3d and Supplementary Table 11). Interestingly, variant rs146239222 is found in an enhancer region^48, 49^ with high H3K27ac modification, DNase clusters, and multiple transcription factor binding, which is also consistent with the output of the CARMEN annotation module (Supplementary Fig. 10c,d), and is associated with the expression of MPHOSPH9 in GTEx (*p* = 1.34 × 10^−91^, ENSG00000051825.10)^50^.

**Figure 3:**
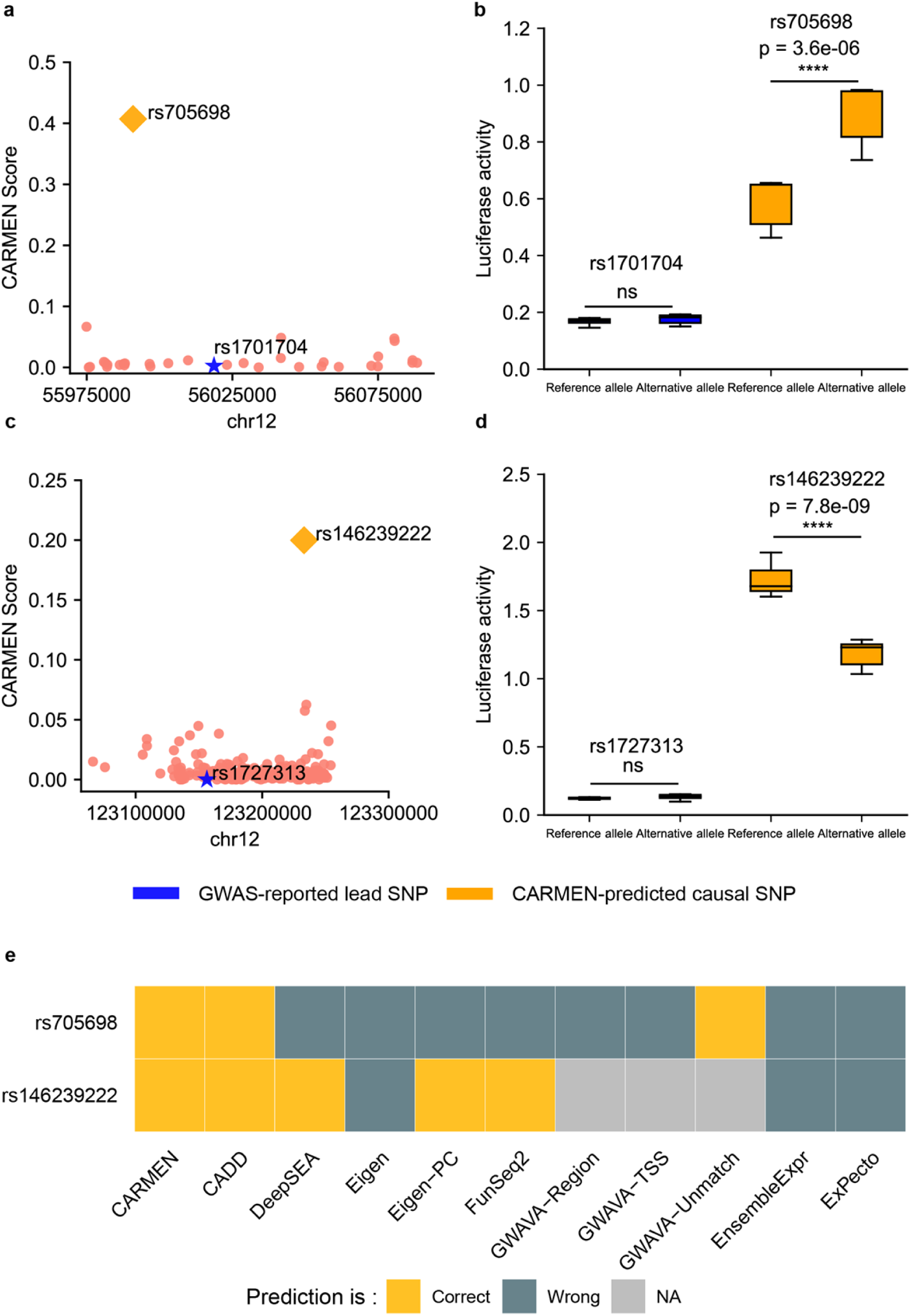
Prioritizing causal variants over linkage variants with lead SNPs in GWAS. **a** and **c,** The X axis represents the genome location of each variant; the Y axis represents the CARMEN score. The blue star represents the lead SNP, which has been reported in the GWAS Catalog. The dots represent the variants with an LD r^2^ greater than 0.75 (CEU) with the lead SNP. The yellow diamond represents the causal variant with best CARMEN score. **b and d**, Luciferase report assays well validated the prediction in type 1 diabetes mellitus (**b**) and type 2 diabetes (**d**). The results were from more than 8 technical replicates with 3 independent experimental replications. **e**, Comparing the performances of CARMEN and other tools on the two luciferase assays that validated as the positive variants above.

## Discussion

Expression quantitative trait loci (eQTL) explain the variant effects on gene expression at the mRNA level^51^. When we applied CARMEN on all 7,627,599 multi-tissue *cis*-eQTLs reported by GTEx v7^52^, we also found several cases where the reported lead SNP showed significantly weak regulation potential compared with that of the linked variants (Supplementary Table 6). For example, the SIK2-correlated variant rs1784782 shows very weak regulatory potential (CARMEN score = 0.0004), while the linked variant rs59921976 presents rather high potential (CARMEN score = 0.6091). The independent MPRA assay confirmed that the CARMEN-predicted causal variant rs59921976 had significant expression changes, but the reported lead SNP rs1784782 did not^53^, and the validated expression-modulating variant rs59921976 was also missed by ExPecto as a nonsignificant variant.

We calculated feature importance for all CARMEN-E and CARMEN-F features. Consistent with previous reports^28, 54^, evolutionary conservation is found to be important for both modules. Furthermore, transcription factor binding profiles play major roles in the CARMEN-E module, while histone markers are more “important” for CARMEN-F, suggesting different aspects of the two modules model (Supplementary Fig. 9, Supplementary Table 4 and Supplementary Table 5).

CARMEN is well tuned for large-scale data, running at ∼10,000 variants per hour on a modern Linux server (Supplementary Fig. 13). CARMEN is available as both a webserver and a standalone package at http://carmen.gao-lab.org. Designed as a one-stop portal, the CARMEN Web server supports not only on-the-fly prediction but also a user-friendly interface for visualizing the results (Supplementary Fig. 12). To help identify the functional mechanism of noncoding variants’ regulation-modulating effects, the Web server provides statistics and specific information about these features in the “Task Results” box as “show details” to help users understand the prediction results.

## Online Methods

### Learning regulation code by sequence-orientated model

#### Predict variants’ effect with sequence-orientated models

The raw data for 1,249 transcription factor (TF) binding profiles, 766 histone markers, 280 DNase I – hypersensitive sites and 108 DNA methylation profiles were downloaded from ENCODE (Supplementary Table 1). We filtered the data to ensure quality. For TF data, only those involved in conservative irreproducible discovery rate (IDR) peaks and optimal IDR peaks were included. For the data about histone marker profiles, replicated peak data were included. For DNase I – hypersensitive sites, pseudoreplicated IDR peak data were used. All data were in bed format, and the reference genome version was GRCh38. The same features (such as CTCF) from different cell lines and using different experimental treatment methods were considered as different chromatin features.

To prepare the input for the model, we took a 200 bp window centered on the ChIP-seq peak or target methylation site from all data. For some features, there were several data files from different labs or experiments, and as a result, these features were merged. The merging standard used was that if two sequences from two data files for the same feature overlapped each other, the sequence with a lower peak value was abandoned. Then, the bed files were converted to FASTA files to obtain sequences from the reference genome. During this process, the reverse complement strands were also converted (Supplementary Fig. 4).

For the classification model, positive datasets and control datasets were both needed. For ChIP-seq data, we removed the positive data from the reference genome and then split the rest into 200 bp bins. We randomly selected some bins to make the number of the positive and control dataset equal. For methylation data, if the methylation rate of the target site was higher than 50%, it was considered positive data; if the rate was lower than 50%, it was considered control data; and if the rate was 50%, it was abandoned. Positive data were labeled as “1” and control data as “0”. Training and testing sets were split randomly in an 85% to 15% ratio. To obtain a reliable and stable model, five-fold cross-validation was conducted.

#### Structure of the model

The classification model was a deep convolutional network (Supplementary Fig. 5). The basic layer types in the model were a convolution layer, a pooling layer and a fully connected layer. The first layer of the model was a 1D-convolutional layer, which was responsible for detecting features along the sequences with the activation function ReLU. This layer had 128 or 256 kernels, and the length of each kernel was 4, 10, 12 or 20. Each kernel could be considered a PWM calculator that moved on the sequence with step one. Formally,

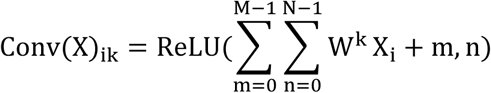

where *X* is the input, *i* is the index of the output position and *k* is the index of the kernel. *W^K^* is the weight matrix whose shape was M × N, *M* is the kernel length and *N* is the channel number of the input. The second layer was a 1D–max-pooling layer with a pooling size of 10 or 20. It was used to take the maximal value from nonoverlapping windows. The function of the max-pooling layer was to reduce the dimensionality of the input data. It was defined as

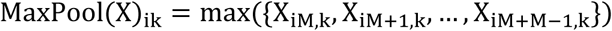

where *M* is the pooling size.

The following layers were the dropout layers and fully connected layers with ReLU as the activation function. The fully connected layer multiplied linear combinations of the input and was defined as:

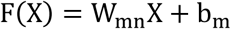

where *m* is the output channel number and *n* is the input channel number. The dropout layer was used to reduce overfitting by randomly setting a certain proportion of input units to zero during training. We set *m* to 512 the drop rate to 0.5.

The last layer of the model was a fully connected layer with a sigmoid function as the activation function. The sigmoid function scales the prediction to 0-1 and is defined as

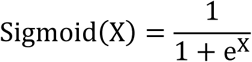

#### Training of the model

The loss function of the model was binary cross entropy (BCE). Specifically,

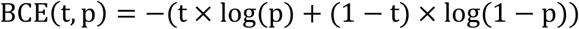

where t is the true label and p is the prediction of the model.

The optimizer was stochastic gradient descent (SGD), and the learning rate was 0.01. Early stop was used during the training process. The model ran 50 epochs for training. The classification model’s performance was evaluated in terms of the classification rate by using the area under the receiver operating characteristic curve (AUROC).

We obtained a 100 bp flanking sequence upstream and a 99 bp flanking sequence downstream of each allele, resulting in a 200 bp reference sequence and a 200 bp alternative sequence for each variant. With each sequence-oriented model, we calculated the log2 fold changes (as the method shown in DeepSEA) of the reference and alternative sequences as the variant effects (Supplementary Fig. 5).

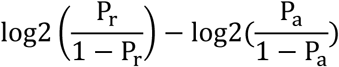

#### Implementation

Implementation of the model utilized the deep learning framework TensorFlow (https://www.tensorflow.org/), and all scripts for the models and evaluations were written in Python (https://www.python.org/). An NVIDIA Tesla P100 Graphics Processing Unit was used to train the model.

#### Predicting variant effects on conformational and physicochemical DNA properties

As Rong Li et al.^22^ showed in their paper, 13 conformational and physicochemical DNA properties are useful to find functional elements in the genome, and thus we used the method referred to by Rong Li et al. to calculate the variant effect on these 13 features.

For the 12 physicochemical features selected from DiProDB^55^, we obtained a 2 bp flanking sequence on both sides of each allele, resulting in a 5 bp reference sequence and a 5 bp alternative sequence. A 2 bp-wide sliding window is used for the 5 bp sequence, and the values from each window are then summed. For the hydroxyl radical cleavage pattern (OH)^56^, we obtained a 6 bp flanking sequence on both sides of each allele, resulting in a 13 bp sequence for each allele. The score was calculated using a 4 bp-wide window for the 13 bp sequence. Because the OH feature was influenced by the double strands of DNA, the OH score for each allele was the average score of the two strands.

#### Predicting variant effects on evolutionary conservation features

Further features include the conservation score of each position of the variants. We used hgwiggle to download 8 kinds of conservation scores: 4 PhastCons and 4 PhyloP scores from the UCSC Genome Browser in reference genome version GRCh37.

#### Feature selection

We annotated each variant with each feature, and we used a multilayer perceptron neural network to introduce the complex relationship of different features to improve the flexibility of feature interactions (Supplementary Fig. 6). The first three layers learn the code to represent input features by the hidden units.

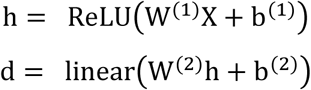

A dense layer with L1 regularization is added behind these layers. We added a λ_l_ = 0.0001 penalty to the model weight and a λ_2_ = 0.001 penalty to the output of this layer to increase the sparsity of features.

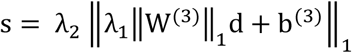

The last two layers of the model are fully connected layers, one with a ReLU activation function and the other with a sigmoid function. The sigmoid function scales the output ranges from 0 to 1.

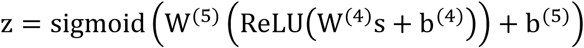

We label the control variants labels as 0 and the positive variants as 1. To solve the binary classification problem of whether the variant has an effect on modulating expression, we trained the multilayer perceptron neural network using 2,424 features with a step size of 300 and then selected the best model as the basic model to perform feature selection.

To learn the important features, we used Deep Learning Important FeaTures (DeepLIFT) ^25^ to calculate the contribution score of each variants’ features by comparing the difference in the activation value of each neuron to its ‘reference activation’. Here, we chose “all zeros” as the reference. After that, we calculated the distribution of the contribution score of the input data and selected the features with contribution scores higher than the absolute cutoff. We selected features with contribution scores greater than 0.5 or less than -0.5 in the CARMEN-E module and contribution scores greater than 2 or less than -2 in the CARMEN-F module. With two training datasets, after feature selection, we then trained the Adaboost Decision Tree with 689 features and the Random Forest with 1,190 features.

The loss function was binary cross entropy, and we used ‘Adadelta’ to optimize the models with default parameters. The batch size was 100, and each model was trained with 100 epochs. We utilized the early-stopping method and set the patience to 30 to avoid overfitting. We used Keras version 1.1.0 and Python version 2.7.12 to implement the model, and NVIDIA-SMI, Driver version 381.22, with a GTX 1080 Ti to train the model.

To analyze the different types of feature importance, scikit-learn 0.19.1 was used to calculate feature importance determined by its contributions to the regulation-modulating variant classification. Then, we calculated the cumulative feature importance for each feature category.

#### Model selection

To optimize the hyperparameters of the model and select the best model for feature selection, we used hidden units, ranging from 1 to the number of features with a step size depending on the number of features. Random-seed for cross-validation, random-seed for models and five-fold cross-validation were used for model training. We then selected the best model after using the AUROC to evaluate the performance of the models.

### Predicting the variant effects model

#### Dataset Curation

In CARMEN prediction, two datasets were used for model training (details in Supplementary Table 7 and Supplementary Table 8). In the expression module, the positive dataset was the mutagenesis data of experimental validated variants, because high-quality labeled variants can facilitate the methods to distinguish regulatory variants. Since 2009, the parallel reporter assay has been used to study promoters^57^, and later, this assay was modified to derive several assays, such as the MPRA and STARR-seq. Here, we curated data from 4 papers that applied MPRA to confirm regulatory variants. In the function module, the positive variants were from HGMD professional version 2018.4 with disease-causing variants in regulatory mutations and the control variants from 1000 Genomes. When comparing the performance of CARMEN-F with other tools, since we randomly selected the 1000 Genomes variants, it is possible that these data were used in training dataset for other tools, and so we curated another independent pathogenic dataset (Supplementary Table 7). We excluded variants that could not map from the GRCh37 to the GRCh38 versions of the human reference genome.

CARMEN prediction consisted of two modules. The first was CARMEN-E, which was trained with an expression-related dataset to evaluate the variants’ effects on gene expression. After the feature selection process, we obtained 689 features. Then, we used the Adaboost Decision Tree to train this dataset with five-fold cross-validation. The other module was CARMEN-F, which evaluated the effect of variants on disease risk. After the feature selection process, we obtained 1,190 features. Random forest with sample weights was used to train the model with a disease-related dataset using five-fold cross-validation. AUROC was used to evaluate model performance. The CARMEN score was calculated with the CARMEN-E and CARMEN-F scores with Bayes’ formula.

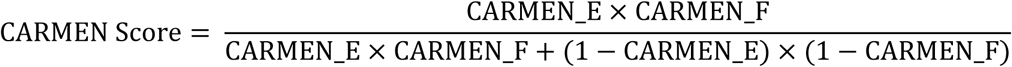

### Model evaluation and application

The model was evaluated with two independent datasets with a large number of variants previously tested by self-transcribing active regulatory region sequencing (STARR-seq) in 2018. The first dataset was curated from a study that systematically identified regulatory variants with cancer risk. In this study, the author identified 1,333 variants in the fragments that regulated gene expression from 10,673 variants. Filtering out the variants’ reference or alternative alleles in the fragments that did not have statistical significance (DESeq2^58^ *p* values) and the locus that could not map from GRCh37 to GRCh38, we obtained 6,539 variants. Among these variants, 1,164 variants in fragments that regulated gene expression and 65 variants had different regulatory activities in two alleles.

We also curated another dataset based on the Biallelic Targeted STARR-seq^59^, which tested 43,500 variants and identified 2,720 variants with significant ASEs. For these two unbalanced datasets, we calculated the weighted accuracy and F1 score to evaluate the performance. The thresholds of the tools that we compared can be found in Supplementary Table 10.

GWAS Catalog v1.0.2 was downloaded from https://www.ebi.ac.uk/gwas/. We also extracted the strongly linked variants (i.e. r^2^ > 0.75) with reported lead SNP based on the haplotypes generated by 1000 Genomes phase 3 across five populations (CEU, CHB, PUR, TSI, YRI). Then, CARMEN was applied to these variants to identify the potential causal variants other than the reported lead SNP. GTEx v7 multi-tissue variants data were from https://storage.googleapis.com/gtex_analysis_v7/multi_tissue_eqtl_data/GTEx_Analysis_v7.metasoft.txt.gz. Metasoft extends the random effects (REs) meta-analysis model into the RE2 model^52^, which was used in GTEx v7 to evaluate the SNP effects on gene expression across multiple tissues. Furthermore, we wanted to identify the variants that were labeled by the RE2 model as lead SNPs but not the causal variants for the eGenes. We used CARMEN to predict all the variants except those without GRCh38 loci or conservation scores. For each gene, the lead SNP was the most significant variant with the smallest *p* value calculated by the RE2 model. We also extracted linkage SNPs with an LD r^2^ greater than 0.75 with each lead SNP in the CEU population. The lead SNPs in the GTEx multi-tissue eQTLs were all predicted by CARMEN. If the lead SNPs were predicted to be negative by CARMEN, then we selected variants with LD r^2^ greater than 0.75 with the lead SNPs. Moreover, we required the high linage variants were not existed in GTEx multi-tissue eQTLs. Then, we used CARMEN to pinpoint the predicted causal SNPs from the negative lead SNPs in each eGene.

### Luciferase reporter assay

For the luciferase reporter assay, HEK293T cells were routinely cultured in DMEM/10% FBS until ready for transfection. Cells were plated in each well of a 6-well plate and transfect the cells at ∼70% confluency with PEI and transfected with 1 µg pGL4.23 firefly vectors containing the selected fragments using standard restriction-enzyme cloning and Renilla plasmids as a transfection control in a 1:1 ratio. Twenty-four hours after transfection, cells were washed twice with cold 1× PBS, and firefly and Renilla luciferase activity was measured using the Dual-Luciferase Reporter Assay System (Promega) according to the manufacturer’s protocol. All the luciferase activity measurements were performed in triplicate for each condition with three independent experimental replicates. Student’s t-test was applied to estimate the statistical significance of the difference in luciferase activity between the two alleles.

### Availability of data and materials

All source codes and data are available freely for academic usage at http://carmen.gao-lab.org. The scripts for generating all figures could be downloaded via http://carmen.gao-lab.org/download/CARMEN-results-source-codes.tar.gz

## Acknowledgments

The authors thank Drs. Zemin Zhang, Cheng Li, Letian Tao, Jian Lu and Liping Wei at Peking University for their helpful comments and suggestions during the study. This work was supported by funds from the National Key Research and Development Program (2016YFC0901603), the China 863 Program (2015AA020108), the State Key Laboratory of Protein and Plant Gene Research and the Beijing Advanced Innovation Center for Genomics (ICG) at Peking University. The research of G.G. was supported in part by the National Program for Support of Top-notch Young Professionals. Part of the analysis was performed on the Computing Platform of the Center for Life Sciences of Peking University and was supported by the High-performance Computing Platform of Peking University.

## Authors’ Contributions

G.G. conceived the study and supervised the research; F.Y.S. and Y.W. designed and implemented the computational framework as well as the website; F.Y.S., Y.W. and Y.L. completed the data curation; H.D., F.Y.S, N.L., and X.W.C. contributed to the independent luciferase reporter validation; F.Y.S. and G.G. wrote the manuscript with comments and input from all coauthors.

## Ethics approval and consent to participate

Not applicable

## Consent for publication

Not applicable

## Supplementary information

Supplementary Tables

Supplementary Figures

## Supplementary Figures

**Supplementary Figure 1.**
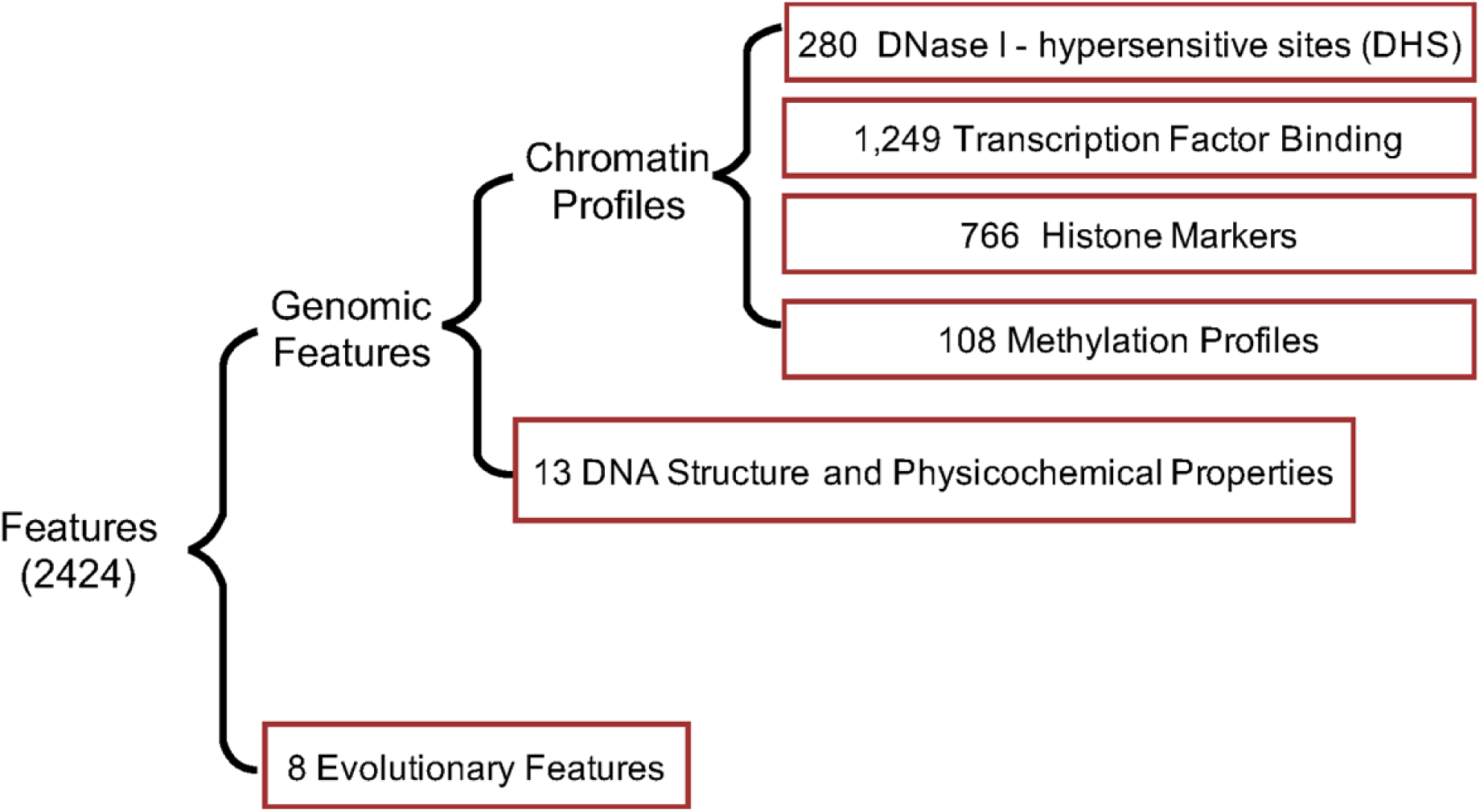
The variant annotations of the CARMEN framework. Each input variant had 2,424 features, the genomic features were all annotated with sequence-based models, and the 8 evolutionary features included PhastCons and PhyloP scores, which were downloaded from the UCSC genome browser.

**Supplementary Figure 2.**
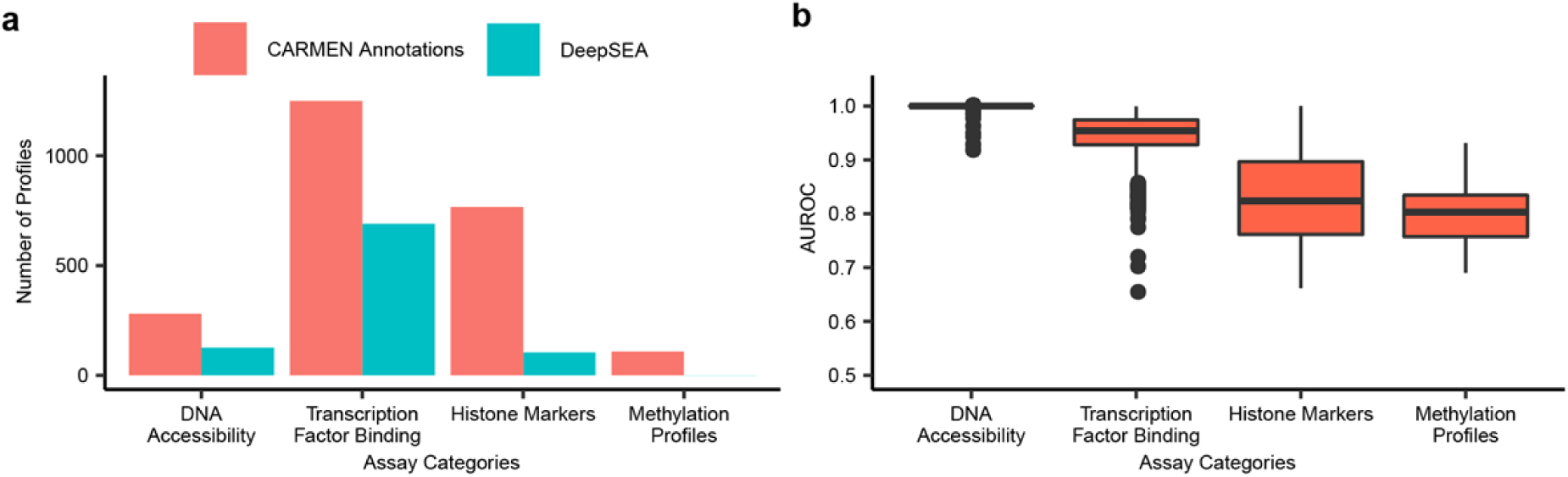
Comparison of the training profiles between the CARMEN annotation module and DeepSEA. (a) The training profiles of the convolutional neural network were the features with 2,403 chromatin profiles from ENCODE, which was greater than the number of training profiles used for DeepSEA (919). Moreover, DeepSEA did not incorporate the DNA methylation profiles. (b) The AUROC of the model performance of the chromatin profile features in CARMEN with 2,403 features; each feature has a separate prediction model, and the performance was evaluated on 15% of training data. The Y axis is the AUROC score.

**Supplementary Figure 3.**
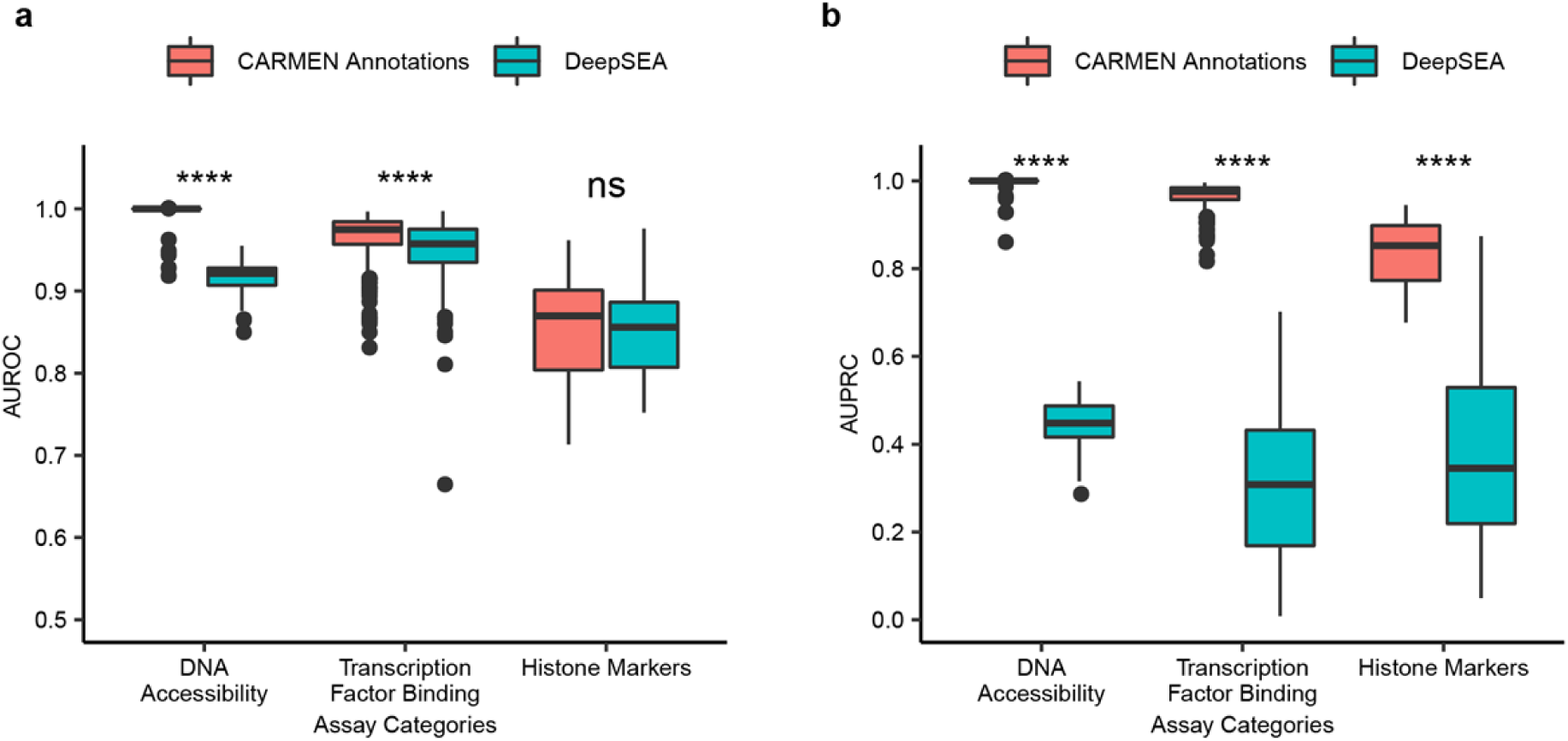
Classification performance of the common features of the CARMEN annotation module and DeepSEA via AUROC (a) and AUPRC (b). There are 466 common features between the CARMEN annotation module and DeepSEA; the performance of the CARMEN common feature prediction models were evaluated on independent testing sets, and the performance of DeepSEA is extracted from the original paper^12^.

**Supplementary Figure 4.**
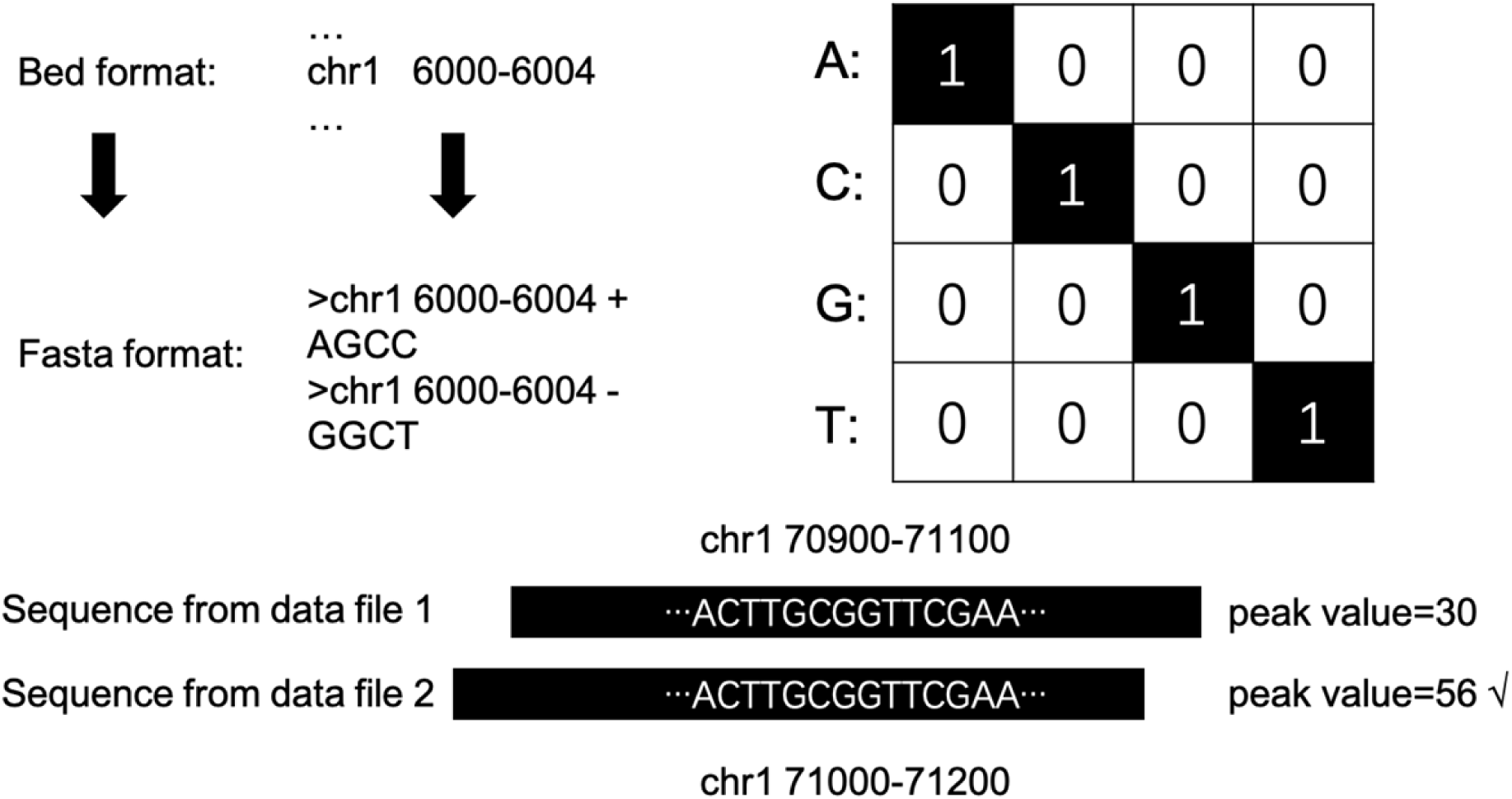
Illustration of data merging, converting to FASTA format and one-hot encoding. Two sequences from two data files overlap each other, and the second sequence has a higher peak value; as a result, it is included in the dataset. Files in bed format are converted to FASTA format. An illustration of one-hot encoding: each sequence is represented by a 200 × 4 binary matrix, with columns corresponding to A, C, G and T.

**Supplementary Figure 5.**
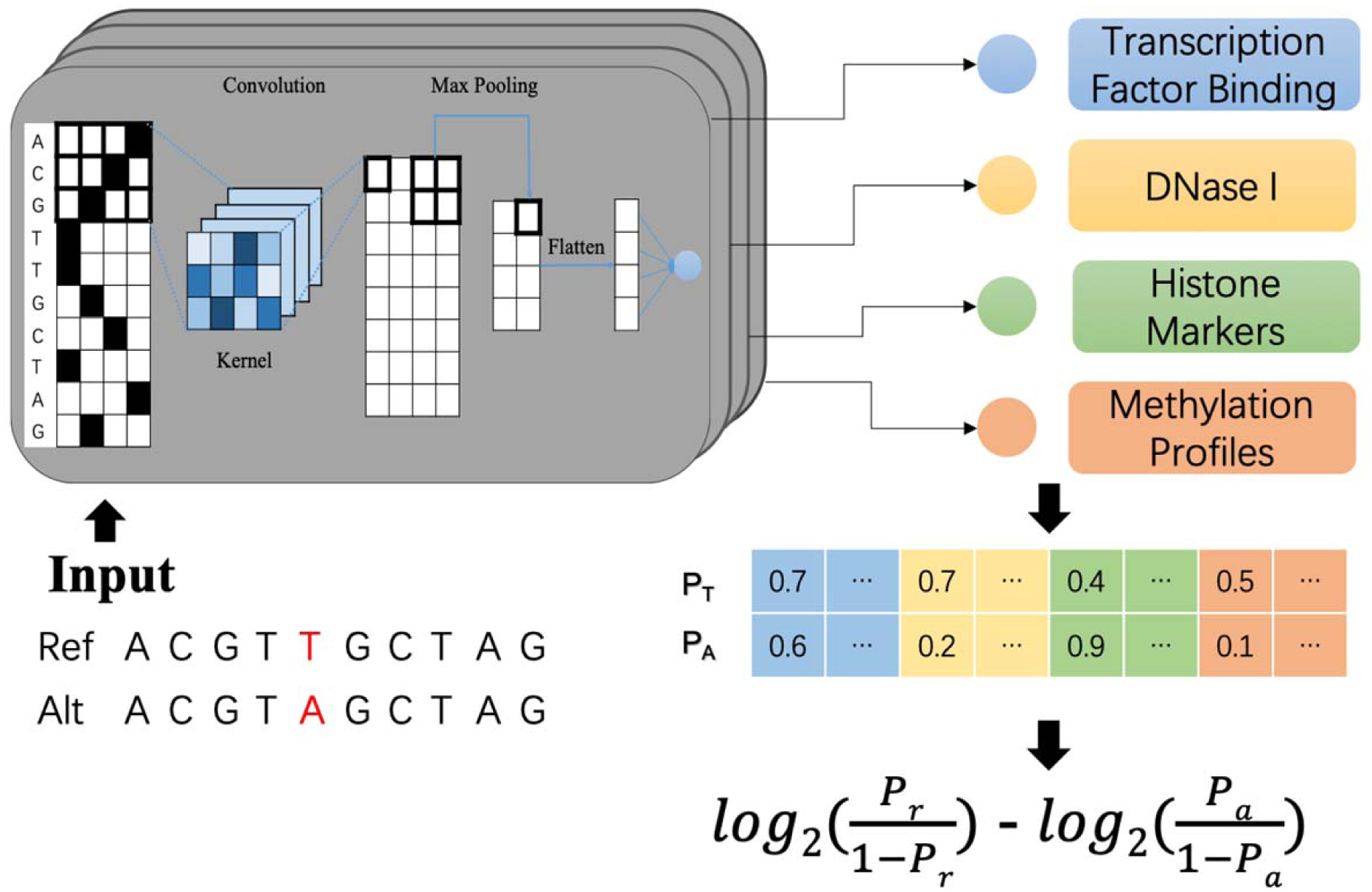
Pipeline for predicting variant effects with sequence-oriented models. The input data are one-hot encoding data. The first layer is a convolution layer for detecting sequence features, and the max-pooling layer reduces the dimensions of the data. Then, after two fully connected layers, a number between 0-1 is generated, which represents the predicted effects in term of log fold change between Ref and Alt sequences.

**Supplementary Figure 6.**
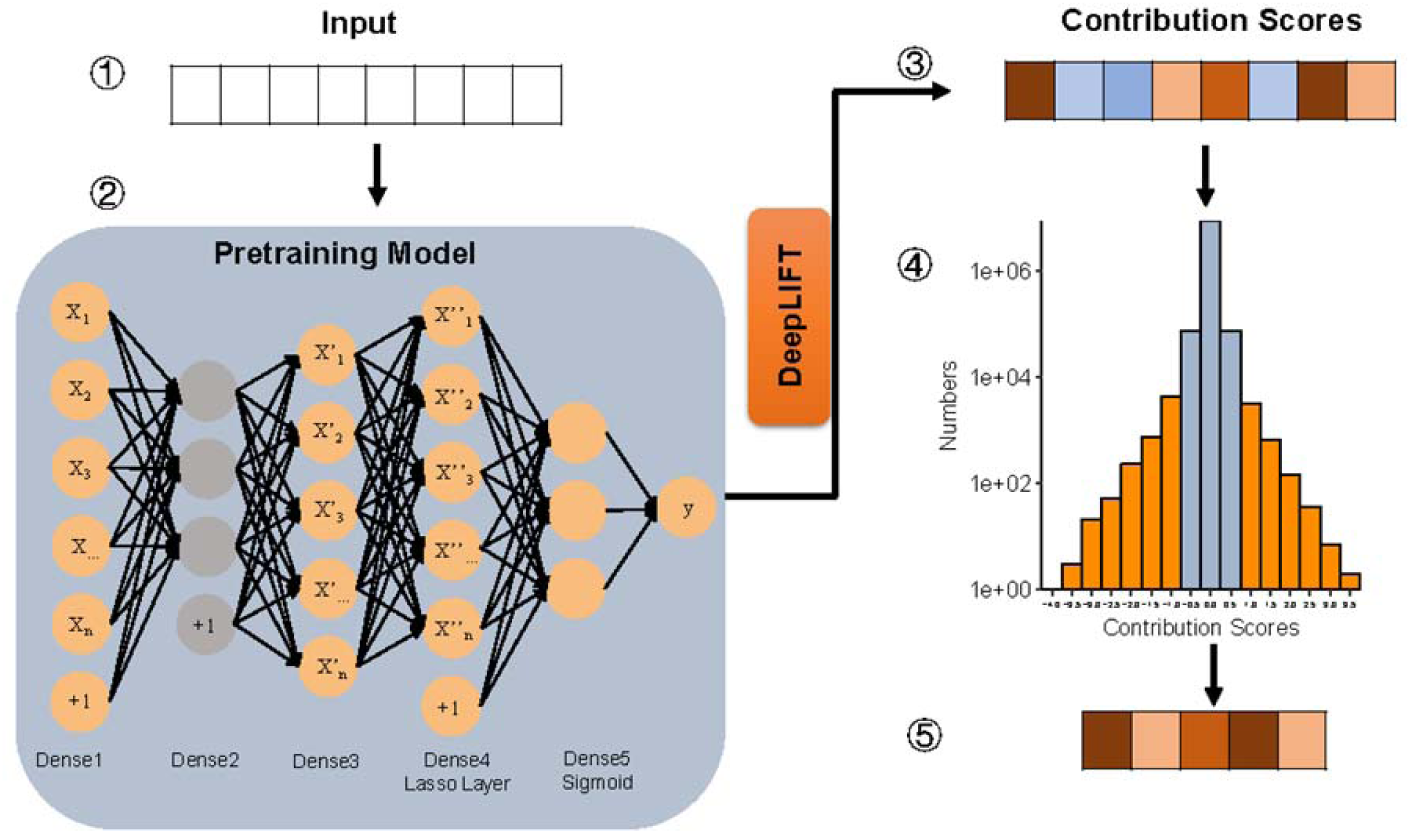
Feature selection procedure. The variants with 2,424 annotated features were used as the input to pretrain a multilayer neural network, then DeepLIFT was used to estimate the contribution scores and obtain the important features for discriminating functional variants.

**Supplementary Figure 7.**
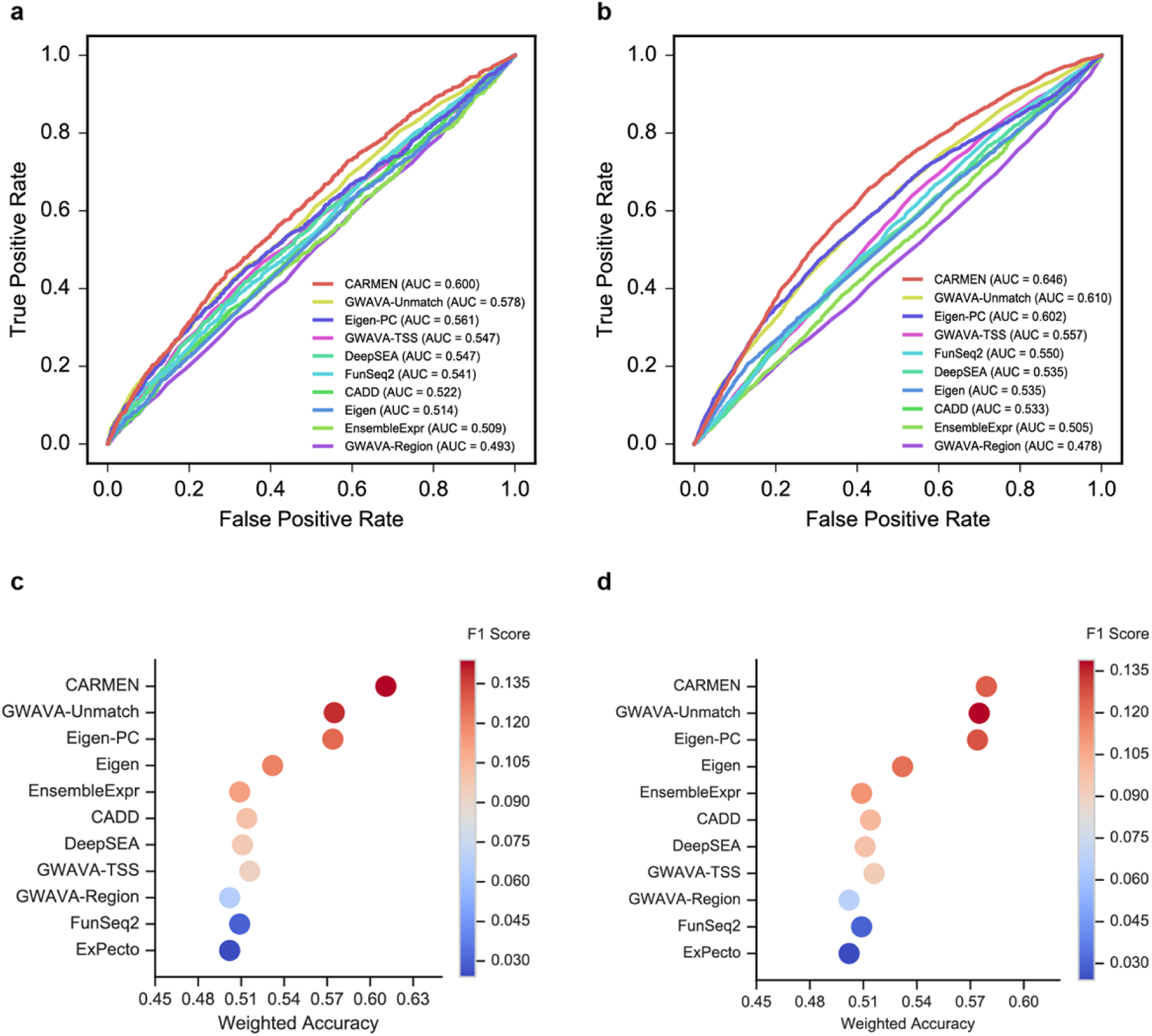
CARMEN performance on a cancer-risk dataset and the BiT-STARR-seq dataset. Comparing CARMEN with other tools to prospect the noncoding-regulation modulating variants. (a) The receiver operating characteristic (ROC) curves for the cancer-risk dataset. Since ExPecto provides the scores of the 208 tissues for each variant, we did not calculate the AUROC and AUPRC of ExPecto. (b) The ROC curves for the BiT-STARR-seq dataset. After curating 43,500 SNPs from Kalita, Cynthia A., et al.^59^ as the second dataset, CARMEN still had the best performance. Among 43,500 variants, 2,720 SNPs with an FDR less than 10% were considered significant ASE variants, and this dataset was called the BiT-STARR-2018 dataset. CARMEN had the highest AUROC (0.646) on this dataset. (c) CARMEN performance using the best AUROC in the BiT-STARR-seq dataset as the threshold. Since this independent testing dataset is imbalanced with a 1:15 ratio of positive to control variants, we calculated the weighted accuracy, which balanced the sensitivity and specificity, and the F1 score, which balanced precision and recall, to compare CARMEN performance to that of the other tools. Although CARMEN had a comparable F1 score with that of the GWAVA Unmatched model, CARMEN had the highest weighted accuracy (0.579) among all tools tested; it performed particularly well in terms of sensitivity. The X axis indicates the weighted accuracy, the Y axis represents different tools, and the bubble color represents the F1 score. The thresholds of different tools were supplied by the official tools’ website or reference papers (ordered by weighted accuracy). (d) CARMEN performance, using the best AUROC in the cancer risk dataset as the threshold.

**Supplementary Figure 8.**
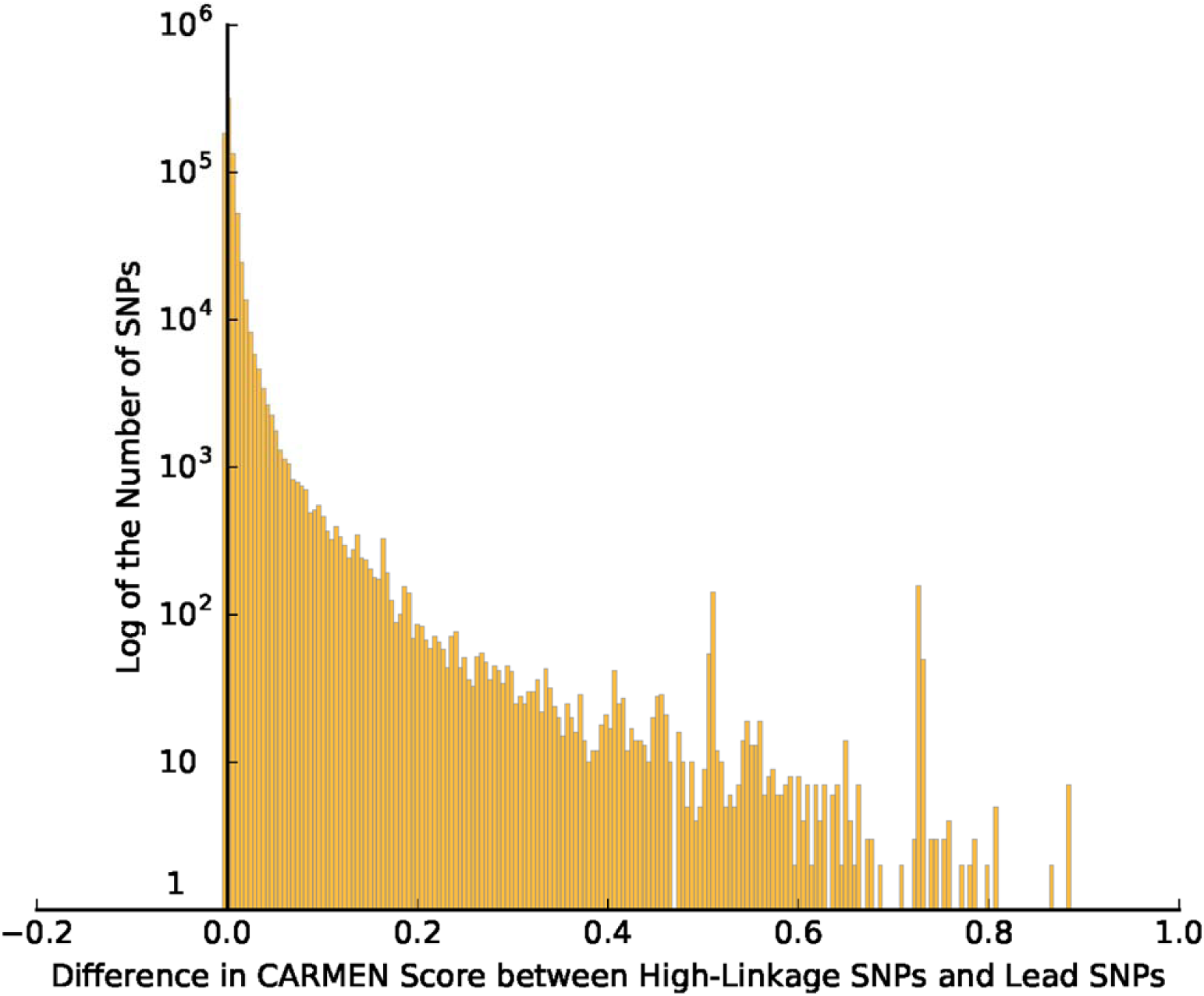
The distribution of GWAS lead SNPs with low CARMEN scores The lead SNPs in the GWAS Catalog, which have CARMEN scores lower than 0.005 in the European, Han-Chinese and African populations, were selected. Then, we calculated the CARMEN score difference between the selected lead SNPs and their high linkage SNPs (r^2^ > 0.75). 45.33% of the selected variants had lower CARMEN scores than LD SNPs with the threshold of 0.005. While most of the differences are modest, 6.65% of variants show differences larger than thirty-fold.

**Supplementary Figure 9.**
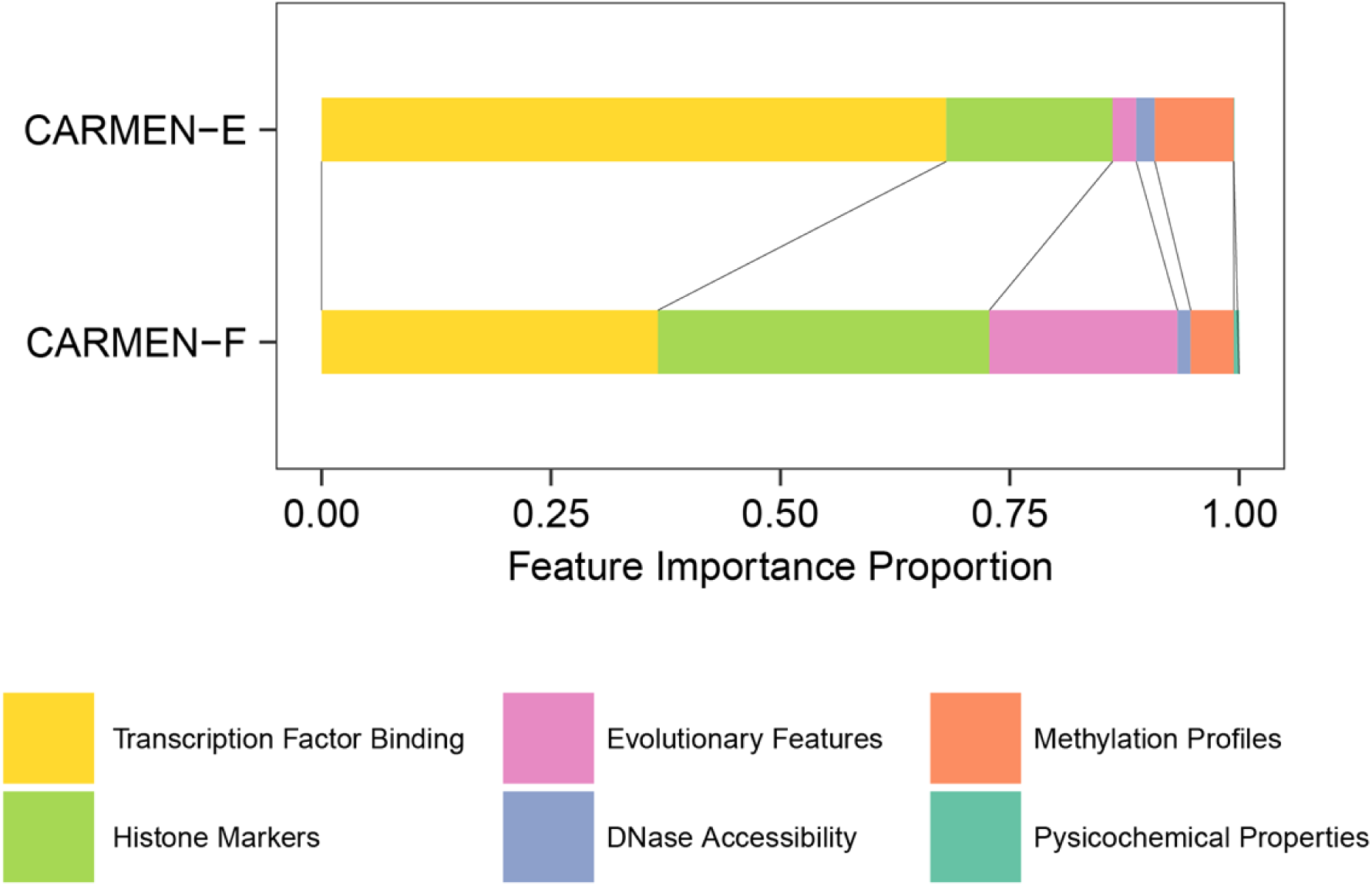
Feature importance proportion. Feature importance was calculated separately in the two modules. The value of the different color bar represents the cumulative importance of each feature category.

**Supplementary Figure 10.**
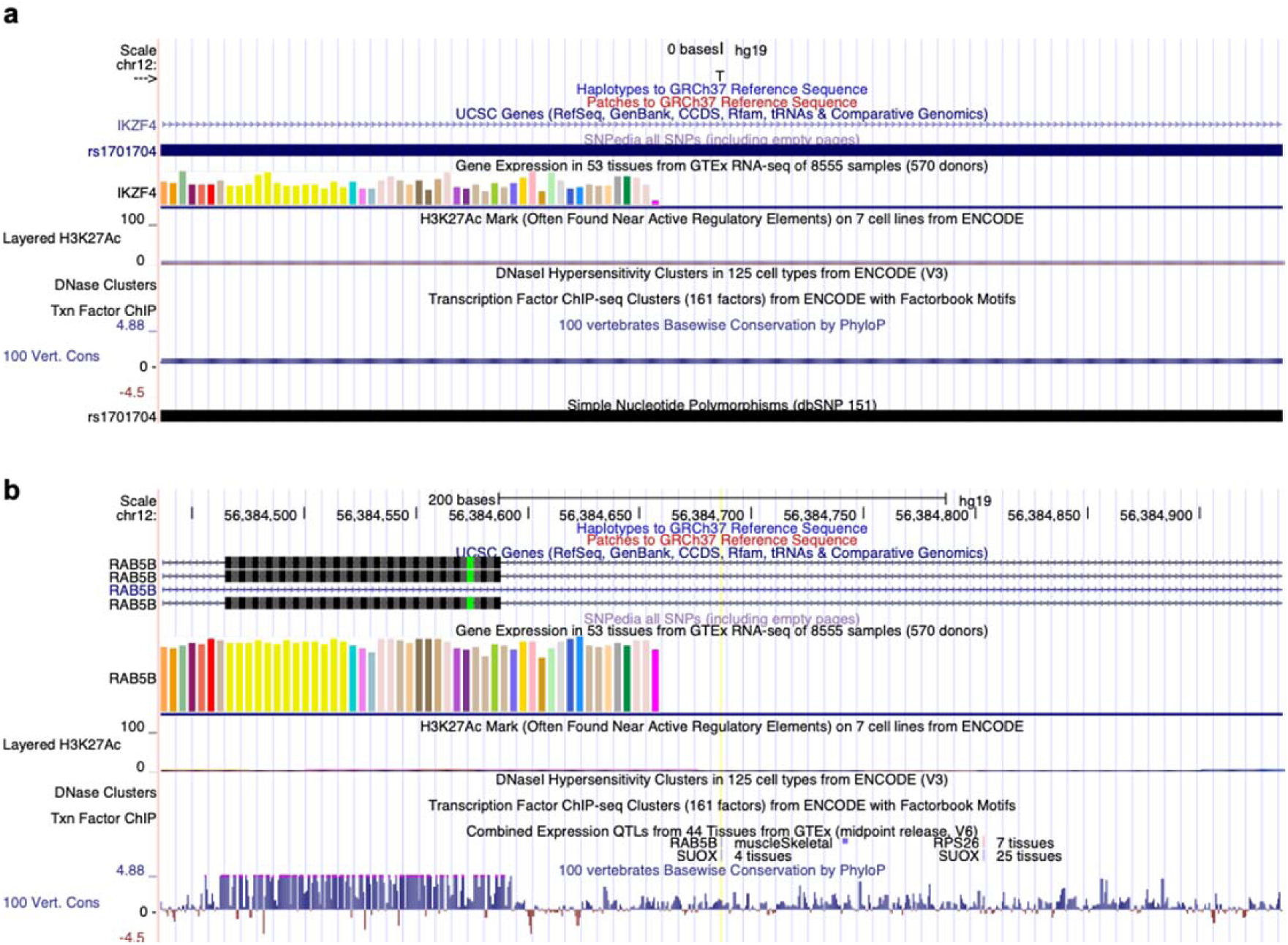
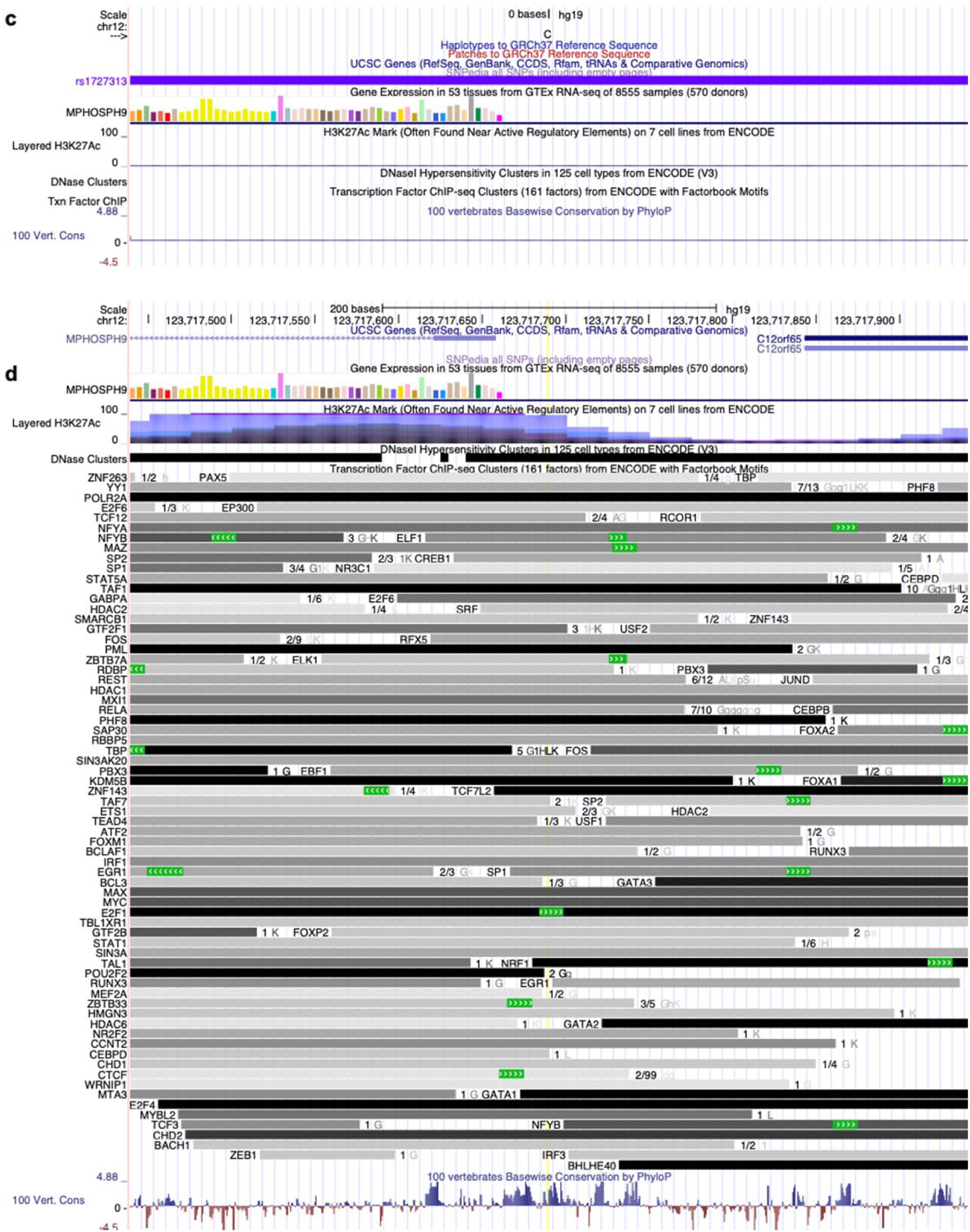
UCSC Genome Browser annotations for variants. (a) Annotations for rs1701704. This variant falls into the intron of gene IKZF4 with low H3K27ac modification but without annotation about DNase cluster and transcription factor binding. (b) Annotations for rs705698. (c) Annotations for rs1727313. (d) Annotations for rs146239222.

**Supplementary Figure 11.**
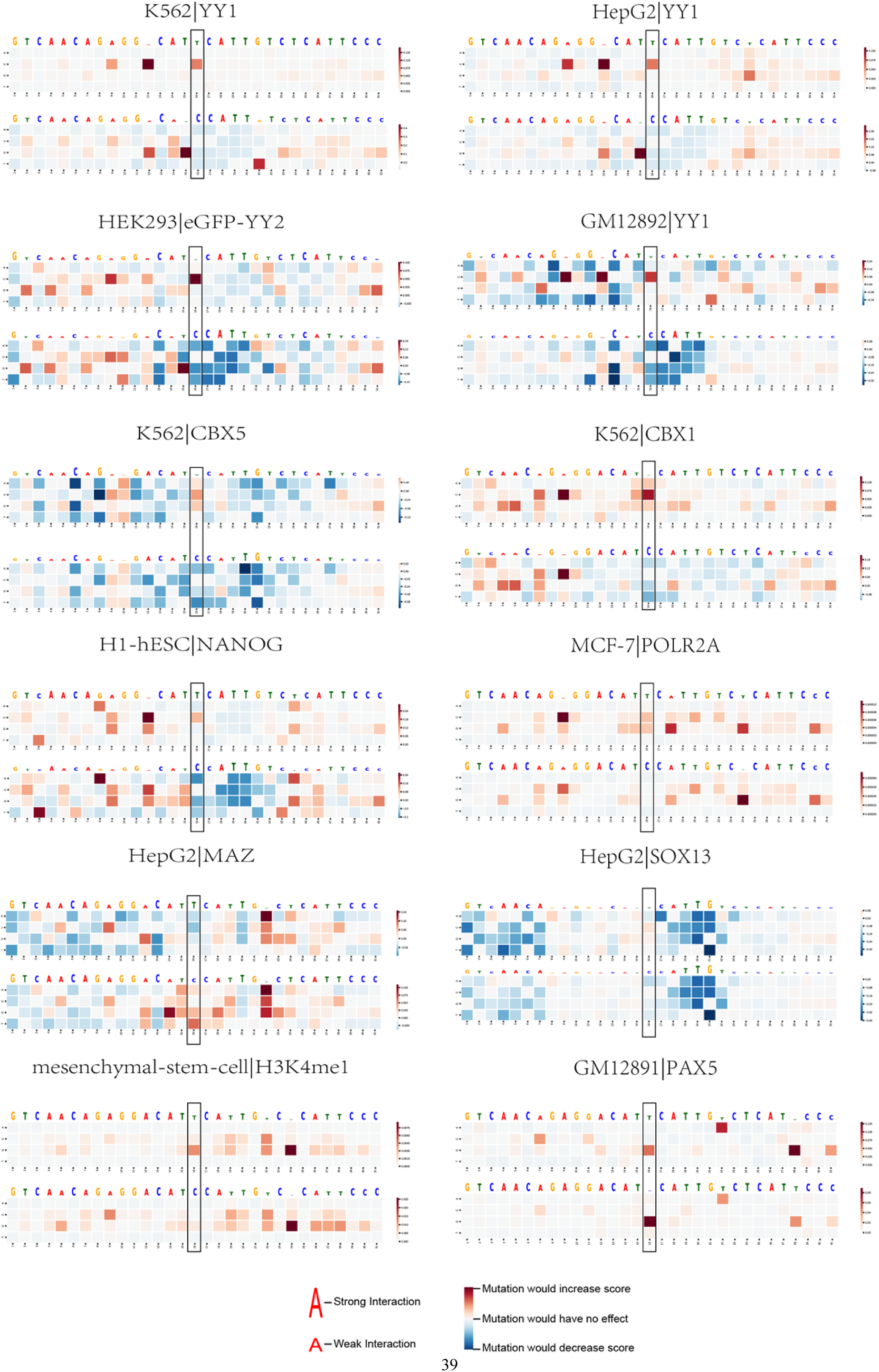
Motifs matched for the sequence around rs705698. Flanking sequences of 15 bp of SNP rs705698 were used to calculate the mutation score. Features annotated by CARMEN with absolute log fold change larger than 1 were selected. The red block indicates that this mutation will increase the binding of this feature. The blue block indicates that this mutation will decrease the binding of this feature. Mutation scores were scaled in each feature. The top heatmap is the mutation score of sequence with reference allele and the bottom heatmap is the mutation score of sequence with alternative allele.

**Supplementary Figure 12.**
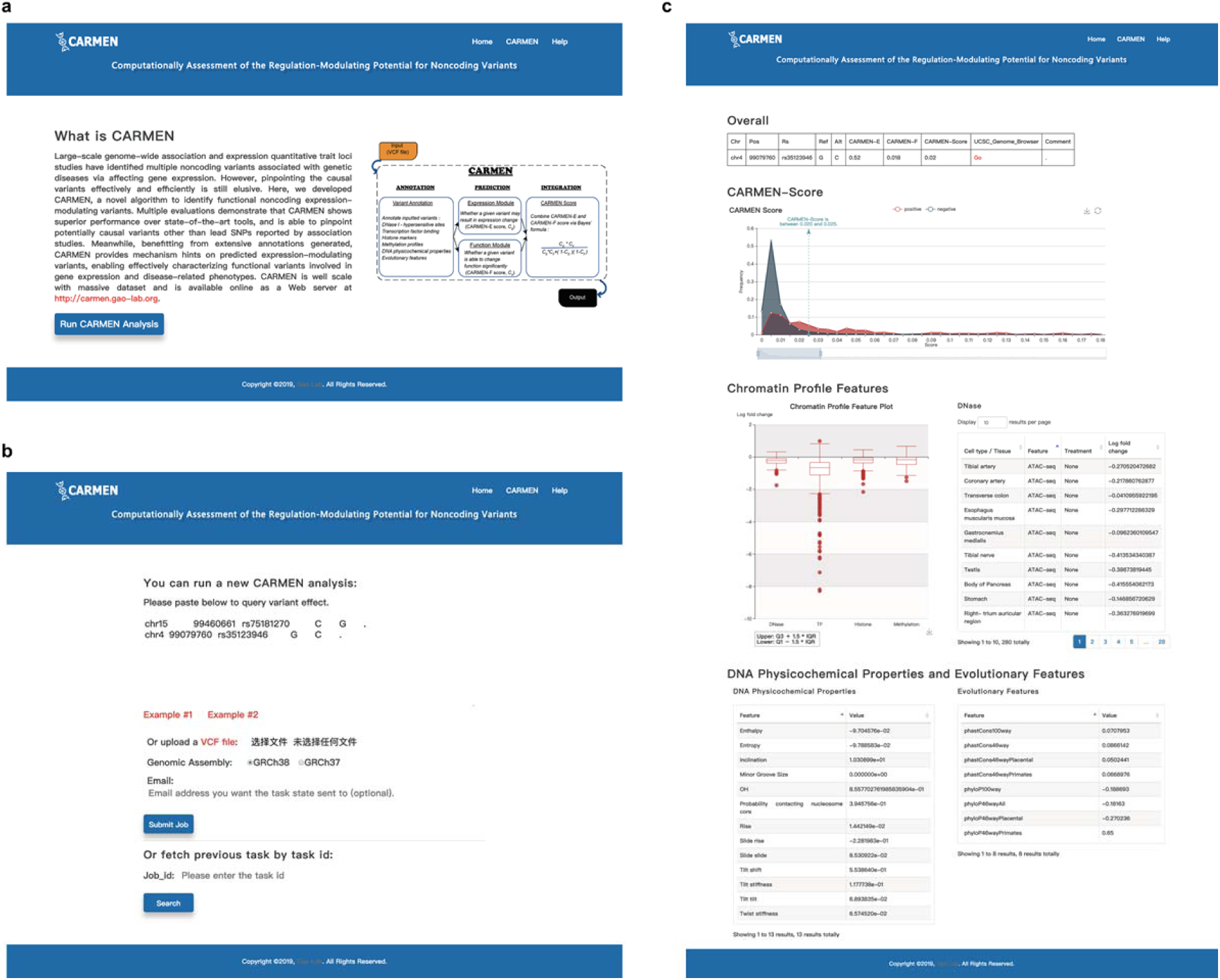
CARMEN Web Server. (a) Home page. (b) CARMEN accepts VCF file as input via both bulk uploading or directly pasting from the browser. (c) Result page with abundant annotations rendered as interactive figures.

**Supplementary Figure 13.**
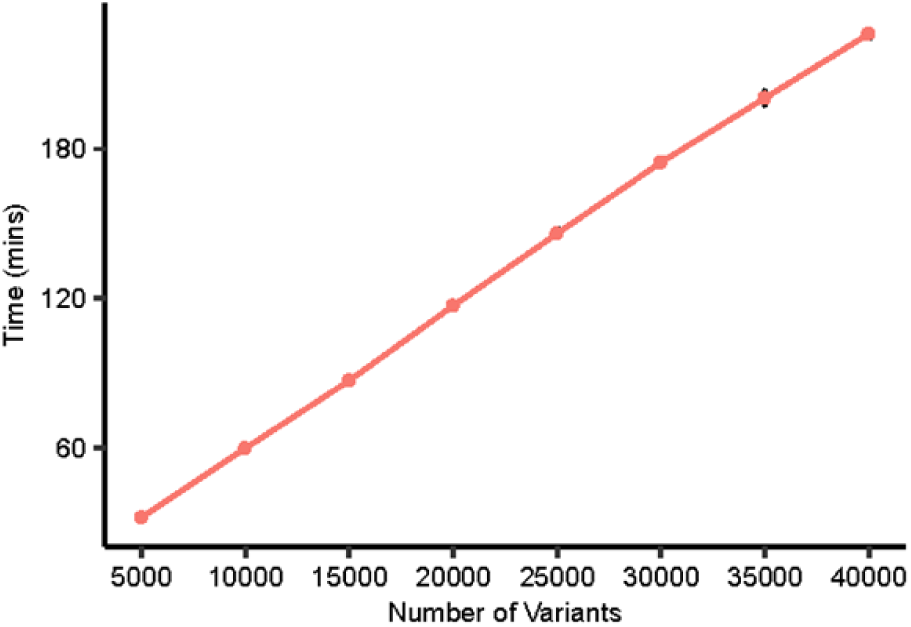
The time cost of CARMEN. Estimating the running time of CARMEN with different number of variants. With 5 times random sampling of different numbers subset from 43,500 variants. Error bar represents mean ±sd.

